# Cdbgtricks: Strategies to update a compacted de Bruijn graph

**DOI:** 10.1101/2024.05.24.595676

**Authors:** Khodor Hannoush, Camille Marchet, Pierre Peterlongo

## Abstract

We propose `Cdbgtricks`, a new method for updating a compacted de Bruijn graph when adding novel sequences, such as full genomes. Our method indexes the graph, enabling to identify in constant time the location (unitig and offset) of any *k*-mer. The update operation that we propose also updates the index. Our results show that `Cdbgtricks` is faster than `Bifrost` and `GGCAT`. We benefit from the index of the graph to provide new functionalities, such as reporting the subgraph that shares a desired percentage of *k*-mers with a query sequence with the ability to query a set of reads. The open-source `Cdbgtricks` software is available at https://github.com/khodor14/Cdbgtricks.

## 1 Introduction

The de Bruijn graph is one of the fundamental data structures that play crucial roles in computational biology. It is tremendously used in various applications, including but not limited to genome assembly [1, 2], read error correction [3, 4], read alignment [5, 6] and read abundance queries [7]. The de Bruijn graph is a data structure in which the nodes represent the distinct substrings of length *k* of a set of strings, called *k*-mers, and the edges link the nodes that share an overlap of (*k* − 1). In a compacted de Bruijn graph, nodes along each maximal non-branching path of the de Bruijn graph are compacted into a single node representing a sequence of length *≥k*, called a unitig. The de Bruijn graph can be constructed from a set of assembled genomes or a set of reads.

Several methods have been proposed in the literature for compacted de Bruijn graph construction, including `PanTools` [8], `Bcalm2` [9], `TwoPaCo` [10], `deGSM` [11], `Bifrost` [12], `Cuttlefish2` [13], `GGCAT` [14], and `FDBG` [15]. As genomic databases grow, there is a demand for dynamic de Bruijn graph data structures supporting sequence additions. `BufBoss` [16], `DynamicBoss` [17], and `FDBG` [15] support the addition and the deletion of nodes and edges in de Bruijn graphs.

Nevertheless, the performance of `BufBoss`, `DynamicBoss` and `FDBG` in sequence addition falls short compared to `Bifrost`. We refer the reader to the results of the `BufBoss` paper [16]. Despite being faster in adding new sequences, `Bifrost` builds a new graph from the sequences to be added, using its construction method, and then it merges the two graphs. The reliance on constructing a graph from the new sequences introduces computational overhead, potentially limiting the scalability and efficiency of `Bifrost`, particularly when dealing with large graphs, highlighting the need for a more efficient updating mechanism. Recent advances have focused on developing memory- and time-efficient indexing structures for *k*-mers. Some notable methods, such as `BLight` [18], `SSHash` [19], `GGCAT` [14], and `Pufferfish` [20], are recognized for their efficiency in this regard. However, it is important to note that these methods are static, meaning that they cannot easily incorporate new data. This limitation becomes especially problematic with large datasets when construction time must be paid for every addition of new sequences.

Indexing methods are used as well for read queries, read mapping, and read alignment on de Bruijn graph. Read mapping against the de Bruijn graph has been studied extensively. `Bifrost` [12], `GGCAT` [14] and `SSHash` [19] proposed read query methods. The limitation of `Bifrost` and `SSHash` is the need to compare any queried *k*-mer with all *k*-mers having the same minimizer, the smallest substring of length *m < k* according to some order. Although it is an efficient design to reduce memory usage, it slows down when negative queried *k*-mers.

In this paper, we propose “`Cdbgtricks`”, a novel strategy, and a method to add sequences to an existing uncolored compacted de Bruijn graph. Our method takes advantage of `kmtricks` [21] that finds in a fast way what *k*-mers are to be added to the graph, and our indexing strategy enables us to determine the part of the graph to be modified while computing the unitigs from these *k*-mers. The index of `Cdbgtricks` is also able to report exact matches between query reads and the graph. We compared `Cdbgtricks` against `Bifrost` and `GGCAT`. Despite `GGCAT` lacking the update feature on the compacted de Bruijn graph, we included it in our comparison due to its potential efficiency in graph construction, which may outperform an update approach. `Cdbgtricks` is up to 2x faster than `Bifrost` on updating a compacted de Bruijn graph on 100 human genomes datasets, and it shows the competitiveness potential against `GGCAT` on larger human genome datasets for which `GGCAT` may not scale due to the high disk requirement. `Cdbgtricks` is up to 3x faster than `Bifrost` and `GGCAT` on updating a compacted de Bruijn graph on a large *E. coli* genomes dataset.

## 2 Methods

### 2.1 Preliminary definitions

A string *s* is a sequence of characters drawn from an alphabet Σ. In this paper, we use the DNA alphabet Σ = {*A, C, G, T*} where every character has its complement in Σ. The complement pairs of Σ are (*A, T*) and (*C, G*). The reverse complement 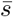 of *s* is found by reversing *s* and then complementing the characters. The canonical string of a string *s* is the lexicographically smallest string between *s* and its reverse complement 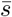. We denote by |*s*| the length of *s. s*[*i*] denotes the *i*^*th*^ symbol of *s*, starting from zero (*s*[0] represents the first character of *s* and *s*[|*s*| − 1] is its last character). Denote by s(*i, j*) the substring of *s* starting at *i* and ending at *j* − 1. A *k*-mer is a string of length *k*. For a given *k*, in our specific context, we denote by *pref* (*s*) = *s*(0, *k* − 1) the (*k* − 1)-prefix of a string *s*, and by *suff* (*s*) = *s*(|*s*| − *k* + 1, |*s*|) the (*k* − 1)-suffix of *s*.

#### Definition 1. de Bruijn Graph

The de Bruijn graph constructed from a set of sequences *S* is a directed graph *G* = (*V, E*) where *V* represents the set of distinct *k*-mers of *S*. When input sequences *S* are made of raw sequencing data, before constructing the graph, the *k*-mers of *S* are counted, and those whose abundance is smaller than a fixed threshold are considered to contain sequencing errors, thus they are discarded. Note that a node *u* represents a *k*-mer *x* and its reverse complement 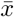. Two nodes *v* and *w* are connected by an edge *e ∈ E* from *v* to *w* if one of the following holds:

1. 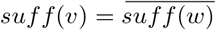
2. 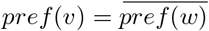
3. *suff*(*v*) = *pref*(*w*)

In any of these three cases, *v* is an in-neighbor of *w*, and *w* is an out-neighbor of *v*. It is worth mentioning that within the scope of this paper, the edges are not stored explicitly; rather, they are deduced from the nodes.

#### Definition 2. Path

A path of a dBG is a an ordered set of nodes where every two consecutive nodes are connected by an edge.

#### Definition 3. Unitig

A maximal non-branching path is a path *p* = {*f, v*_1_, *v*_2_, …, *v*_|*p*| −2_, *l*} where every *v*_*i*_ has only one in-neighbor and one out-neighbor on *p* and *f* and *l* do not have this property. A maximal non branching path can be compacted to form a unitig *u*. The compaction of two nodes *v* and *w* can be achieved as follows:

1. If *suff*(*v*) = pref(w) then *compaction*(*v,w*) = *v* ⊙ *w*[|*w*| − 1]
2. If 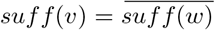 then 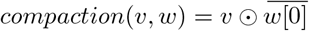
3. If 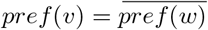then 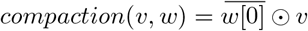

Without loss of generality, we suppose in what follows that two nodes *v* and *w* are in forward-forward direction i.e., *suff* (*v*) = *pref* (*w*). It should be noted that the rest of the cases remain valid within the scope of this definition. In what follows we will not emphasize if a *k*-mer *x* is canonical or not.

#### Definition 4. Compacted de Bruijn Graph

Replacing the maximal non-branching paths by their unitigs provides a compacted form of the de Bruijn graph. An illustration of a de Bruijn graph and its compacted version is shown in Figure 1.

**Figure 1.**
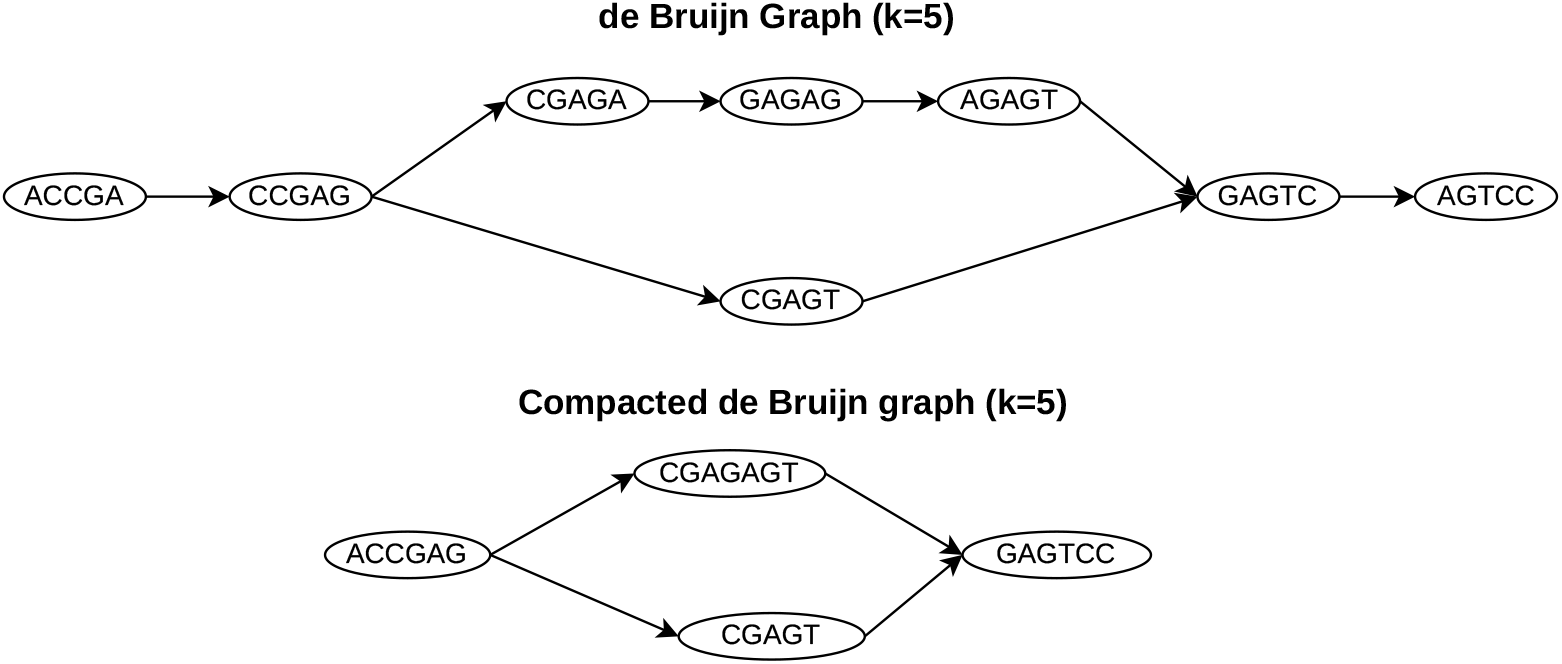
A de Bruijn graph and its compacted de Bruijn graph version.

#### Definition 5. Minimizer

A minimizer of a string *s* is a substring *q* of fixed length *m* where *m <* |*s*| and *q* is the smallest *m*-mer of *s* with respect to some order. In this paper, the order is defined on the basis of a hash function.

#### Definition 6. Minimal perfect hash function MPHF

Given a set of keys *K*, a minimal perfect hash function (MPHF) is a function that bijectively maps the elements of *K* to the elements of the set *I* = {*i*|0 *≤ i <* |*K*|}.

### 2.2 Overview of the Algorithm

`Cdbgtricks` enables to add a set of new sequences *S* to a compacted de Bruijn graph *G*. We denote by *K*_*S*_ the set of *k*-mers in *S* and by *K*_*G*_ the set of *k*-mers in *G*. The set of *k*-mers in *K*_*S*_ but not in *K*_*G*_ have to be added to *G*. We call *N* this set *K*_*S*_ \ *K*_*G*_. To efficiently determine *N* we use the `kmtricks` [21] tool. To help understand the proposed algorithm, we first describe the process when adding *k*-mers from *N* one after another in the compacted de Bruijn graph *G*. We show later (Section 2.4), how to avoid these |*N* | individual additions.

When adding a *k*-mer *x* from *N* to the compacted de Bruijn graph *G*, we distinguish the following cases, as represented in Figure 2:

**Figure 2.**
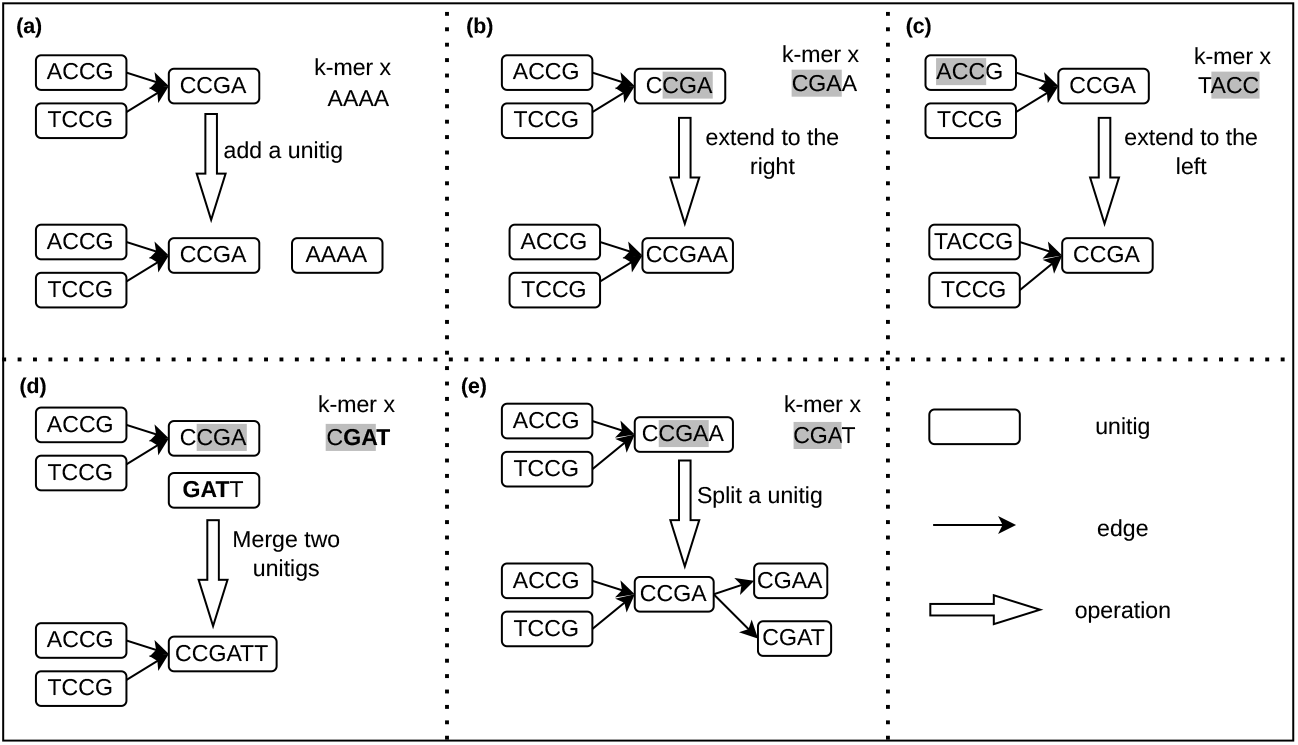
Possible operations when adding a *k*-mer to a compacted de Bruijn graph with. *k* = 4. **(a)** Adding the *k*-mer as a new unitig. **(b)** Extending a unitig to the right. **(c)** Extending a unitig to the left. **(d)** Merging two unitigs. **(e)** Split a unitig into two unitigs. Gray and bold sequences represent overlap between the added *k*-mer and some unitigs of the graph.

1. **Add** *x* **as a new unitig**. Neither *pref* (*x*) nor *suff* (*x*) appears in any unitig of *G*. In this case, the existing unitigs of *G* are not modified and *x* is added as a new single unitig in *G* (Fig 2.a).
2. **Right extension of a unitig**. If *pref* (*x*) equals *suff* (*u*) for a unitig *u* of *G*, and *u* has no out-neighbor, then *u* is extended with the last character of *x* (*u* = *u* ⊙ *x*[*k* − 1])(Fig 2.b).
3. **Left extension of a unitig**. If *suff* (*x*) equals *pref* (*u*) for a unitig *u* of *G* and *u* has no in-neighbor, then the first character of *x* is added to the left of *u* (*u* = *x*[0] ⊙ *u*) (Fig 2.c).
4. **Merge two unitigs**. If the addition of *x* leads to the right extension of a unitig *u*_1_ **and** to the left extension of a unitig *u*_2_, then, after the extensions, *suff* (*u*_1_) = *pref* (*u*_2_). In this case the two unitigs *u*_1_ and *u*_2_ are merged into a unique unitig *u* = *u*_1_ ⊙ *u*_2_(*k*, |*u*_2_|) (Fig 2.d).
5. **Splitting a unitig**. If *pref* (*x*) exists in a unitig *u* of *G*, not being the suffix nor the prefix of *u*, then *u* is split into two unitigs *u*_1_ and *u*_2_ where *suff* (*u*_1_) = *pref* (*u*_2_) = *pref* (*x*) and *x* is added as a single unitig. Respectively, if *suff* (*x*) exists in a unitig *u* of *G*, not being the suffix nor the prefix of *u*, then *u* is split into two unitigs *u*_1_ and *u*_2_ where *suff* (*u*_1_) = *pref* (*u*_2_) = *suff* (*x*) and *x* is added as a single unitig (Fig 2.e).

These operations rely extensively on the pattern matching of *suff* (*x*) and *pref* (*x*) in the unitigs of *G*. In order to rapidly perform these operations, we propose to index the graph, as explained in the next section.

### 2.3 Indexing the graph

A core operation in `Cdbgtricks` consists in identifying if a (*k* − 1)-mer occurs in any unitig of a compacted de Bruijn graph *G*, and, if this is the case, to determine the couple(s) (unitig id, offset) where it occurs.

This operation is performed twice for each *k*-mer *x* of *N* (for *pref* (*x*) and *suff* (*x*)). It has to have a *O*(1) time-complexity and to be fast in practice. This is a very common operation for which existing indexing solutions such as [18, 19] are convenient. However, in the context of this work, the specificity is that, when adding sequences to the graph, the indexed data evolve as some unitigs *G* can be split, merged, extended, and some new unitigs can be added to *G*. Hence, those static methods are not adapted. We propose the following strategy to cope with this particular situation.

#### 2.3.1 Indexing *k*-mers for querying (*k* − 1)-mers

Despite the fact that we query (*k* − 1)-mers, we chose to index *k*-mers instead of (*k* − 1)-mers. A (*k* − 1)-mer may have up to eight occurrences in the *G* because it can be the suffix of four possible *k*-mers and the prefix of four possible *k*-mers. Indexing from one to eight couples (unitig id, offset) per indexed element is not efficient as it requires a structure of undefined and variable size. This leads to heavy data-structures and cache-misses on construction and query times. To cope with this issue, we chose to index *k*-mers instead of (*k* − 1)-mers. Indeed, each *k*-mer of a compacted de Bruijn graph, occurs at exactly one couple (unitig id, offset).

Given this indexing scheme in which *k*-mers are indexed, when querying a (*k* − 1)-mer *x*^*′*^, the eight possible *k*-mers containing this (*k* − 1)-mer (four *k*-mers in which *x*^*′*^ is the prefix, and four *k*-mers in which *x*^*′*^ is the suffix) are queried. If a match is found, the offset of the (*k* − 1)-mer is deduced depending on the case (either *x*^*′*^ is the prefix or the suffix of a queried *k*-mer for which a match is found).

As a matter of fact, we only index each *k*-mer in its canonical form. Then, a queried *k*-mer is searched in its canonical form.

#### 2.3.2 Partitioning the *k*-mers of the graph

Conceptually, we could use any associative table such as a hash table for mapping each *k*-mer of *G* to its couple (unitig id, offset). However, this would require explicitly storing the *k*-mers which is a waste of space as *k*-mers are already explicitly existing in unitigs. Alternatively, we use an MPHF *f* from the *k*-mers of the graph. Doing so, we need only to store the position of each indexed *k*-mer. Formally the position of a *k*-mer *x* is defined by *p*_*x*_ =*< u*_*id*_, *u*_*off*_, *orientation >*, where *u*_*id*_ is the identifier of the unitig *u*, 0 *≤ u*_*off*_ *≤* |*u*| − *k* is the offset of *x* in *u*, and *orientation* is a boolean variable that is true if *x* is in its canonical form in *u*, else it is false. The positions of the *k*-mers in the graph are stored in a vector *V*. Given a *k*-mer *x*, its position is *p*_*x*_ = *V* [*j*] where *j* = *f* (*canonical*(*x*)). There are two observations to be made here:

- At query time, *f* can give valid hash values for alien *k*-mers, which are *k*-mers that are not present in the graph. To handle this, we compare the queried *k*-mer to the actual *k*-mer in the graph, whose position is retrieved thanks to *V* [*f* (*canonical*(*x*))].
- Adding new *k*-mers requires to recompute *f*.

This last point is problematic, since for every addition operation, *f* must be recomputed, which is a linear-time operation in terms of the number of indexed *k*-mers. To resolve this, we divide the set of *k*-mers in the graph into multiple subsets called “buckets”. Each subset is indexed using its own MPHF. The key idea being that while adding sequences to a graph, only a subset of the buckets are modified, and so only a subset of the MPHFs have to be recomputed. At query time, the bucket of the queried *k*-mer *x* is retrieved, and the corresponding MPHF provides the position of *x* in the graph.

Formally, we define {*b*_0_, *b*_1_, …, *b*_*n*−1_} buckets. For each of these buckets *b*_*i*_, an MPHF *f*_*i*_ is computed on its *k*-mers. MPHFs are computed using `PHOBIC` [22] as it provides the fastest lookup compared to the state of the art tools that compute MPHFs.

The *k*-mers in the graph are separated into buckets based on their minimizers. The *k*-mers sharing the same minimizer cannot be distributed into different buckets. However, this strategy may result in a well-known problem of non-uniform distribution of the *k*-mers in the buckets [23]. Some buckets could be orders of magnitudes larger than some others. Also, the small buckets are problematic for the construction of an MPHF using `PHOBIC`, as higher number of *bits/*k−*mer* is required for small buckets (see Figure 3). In fact all the methods that construct an MPHF from a small set of keys uses higher *bits/*k−*mer* compared to constructing the MPHF from a larger set of keys.

**Figure 3.**
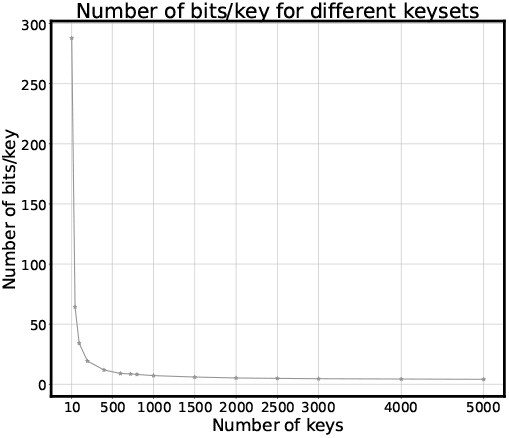
The number of bits/key required for building a MPHF with PHOBIC. MPHFs from different sets of keys of sizes ranging from 10 to 5000 random 64-bit keys were computed by `PHOBIC`, and bits/key were then measured

All in all, we propose a strategy so that all the batches contain a minimum number of *k*-mers.

- The number of *k*-mers sharing the same minimizer should be at least equal to a parameter *ρ* for creating a bucket. Note that the size of a bucket does not have an upper-bound.
- For the remaining *k*-mers, we process them by groups of *k*-mers where the *k*-mers within a group share the same minimizer. From these groups, we create what so-called “super-buckets” which are buckets containing *k*-mers that share different minimizers. We start with an empty super-bucket *S*_0_ to which we add the groups of *k*-mers one by one. Once the number of *k*-mers added to *S*_0_ exceeds *γ* × *ρ k*-mers (with *γ* a user defined multiplicative factor), we create a new super-bucket. We keep creating and filling super-buckets until all groups of *k*-mers are processed. The rationale behind this strategy is to achieve a balanced distribution of *k*-mers on the super-buckets. It is important to note that the size of a super-bucket has an upper bound, which we address in section 2.5.

Finally the data-structure is composed of the following components, represented in Figure 4:

**Figure 4.**
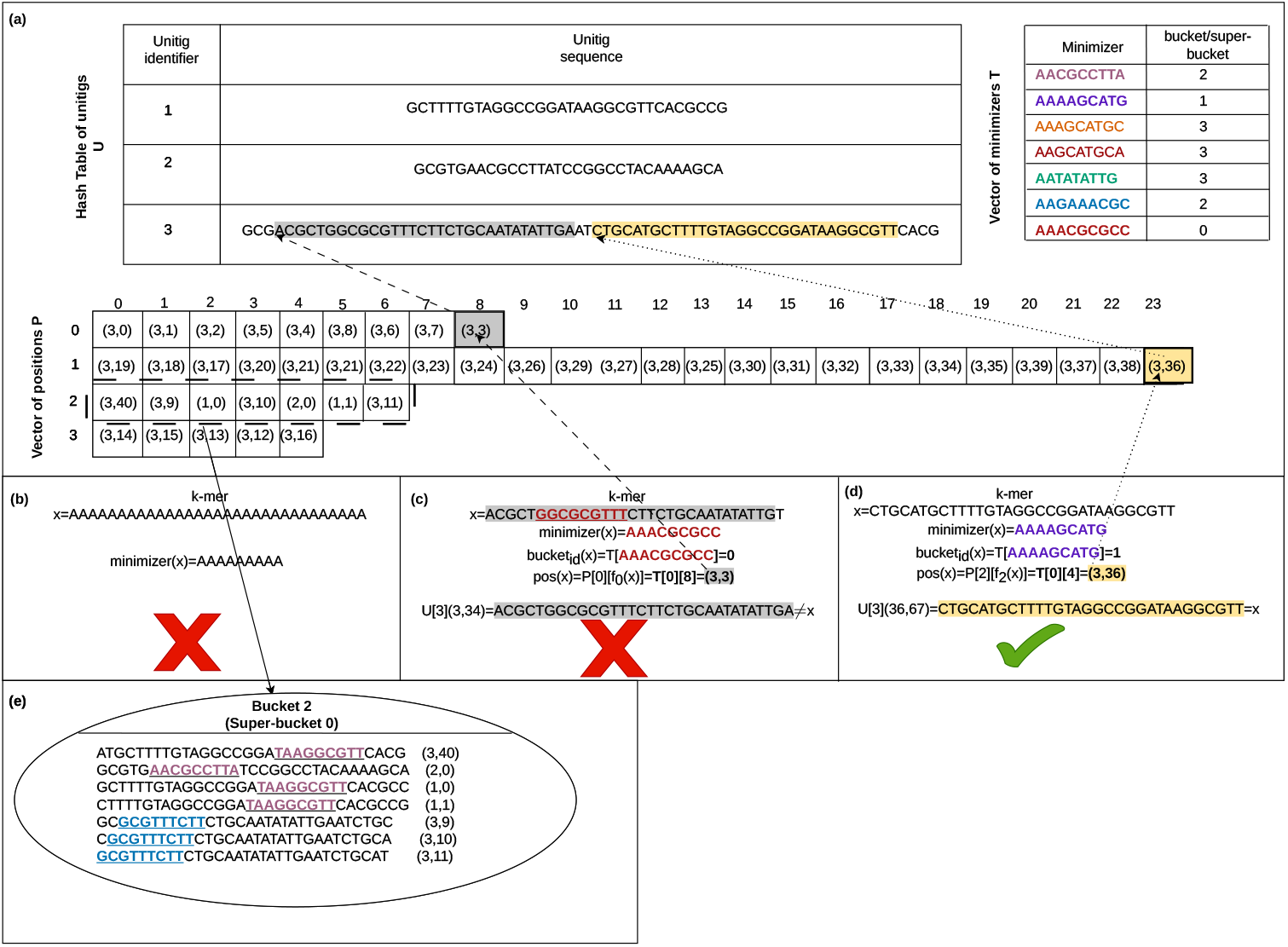
Overview of the data structure. **(a)** the hash table *U* of the unitigs of a compacted de Bruijn graph with *k* = 31; the hash table *T* of minimizers that maps a minimizer to its corresponding bucket or super-bucket; and the vector *P* of positions of the *k*-mers as a tuple (*u*_*id*_, *u*_*off*_) where *u*_*id*_ is the identifier of the unitig in which the *k*-mer occur at offset *u*_*off*_. Note that the values on top of the vector *P* represent the hash value of the *k*-mers computed by the MPHF of their bucket or super-bucket, and the values to the left of *P* represent the bucket or super-bucket identifier. **(b)** querying an absent *k*-mer whose minimizer is not in the index. **(c)** querying an absent *k*-mer whose minimizer is in the index. **(d)** querying a present *k*-mer. **(e)** the dashed box is converted back to a super-bucket of the *k*-mers.

1. A hash table *T* that maps each minimizer to its bucket identifier. The hash table will be used to identify the bucket that may contain a given *k*-mer *x*. The identifier of the bucket is then *b*_*i*_ = *T* [*minimizer*(*x*)].
2. A hash table *F* that maps each bucket identifier to its MPHFs. Hence, *F* [*b*_*i*_] is the MPHF computed from the set of *k*-mers in bucket *b*_*i*_.
3. A hash table *U* that maps the identifier of each unitig of the graph to its sequence. Hence, *U* [*u*_*i*_] is the unitig sequence whose identifier is *u*_*i*_.
4. The positions of the *k*-mers in the graph are stored in a 2-D vector *P*. *P* [*b*_*i*_] is the vector of positions for the *k*-mers in bucket *b*_*i*_. *P* [*b*_*i*_][*F* [*b*_*i*_](*x*)] is the tuple position *< u*_*id*_, *u*_*off*_, *orientation >* for the *k*-mer *x* in the bucket *b*_*i*_.

Overall given the position of a *k*-mer *x, x* can be retrieved by retrieving the unitig *u* = *U* [*u*_*id*_] from the hash table *U*, hence *x* = *u*(*u*_*off*_, *u*_*off*_ + *k*) (Figure 4.c). Note that if the minimizer of *x* is not present in *T*, then *x* does not belong to the graph (Figure 4.b).

### 2.4 Computing the future unitigs and updating the graph

Recall that *N* denotes the set of *k*-mers to be added to a compacted de Bruijn graph *G*. In Section 2.2 we proposed an overview of algorithms in which *k*-mers from *N* are added one after another to *G*. In practice, for performance reasons, we first compact *k*-mers from *N* into what we call “funitigs” (for *future unitigs*).

The funitigs are not simply the unitigs of *N* as any (*k* − 1)-mer of those funitigs that is already in *G* must be either a prefix or a suffix of a funitig. Doing so, the funitigs are not split latter when added to the graph. The details about the funitig construction are given in Algorithm 2 of supplementary materials.

Once the funitigs are constructed, each of them is added to the graph one after the other. The rules described section 2.2 for adding a *k*-mer to *G* exactly apply for adding a funitig to *G*. The `Cdbgtricks` tool exactly implements those rules, that we do not recall here.

### 2.5 Updating the index

`Cdbgtricks` enables to update the index of a compacted de Bruijn graph, after the addition of sequences. The updated index can serve for any future update on the graph and for *k*-mer queries against the graph.

While adding funitigs to the graph, we remember the identifiers of the modified unitigs and of the modified buckets. After all funitigs are added to the graph, the index of the corresponding unitigs and buckets are updated. The update of the index of the graph is tackled into three stages:

1. Splitting a unitig or a joining two unitigs or a unitig with a funitig result in changing unitig identifier(s) and the offsets of some *k*-mers. When we split a unitig *u* into two unitigs *u*_1_ and *u*_2_, *u*_1_ get the identifier of *u, u*_2_ gets a new identifier and the offsets of its *k*-mers get recomputed. When we merge a funitig with one or two unitigs, the resultant sequence gets the identifier of one of these unitigs, and the offsets its *k*-mers get recomputed.
2. The addition of *k*-mers to super-buckets may lead to doubling their maximum size (*γ × ρ*). In this case, for each concerned super-bucket, it is divided into two new super-buckets.
3. Recompute the MPHFs of the buckets and super-buckets to which new *k*-mers were added.

When dividing a super-bucket into two super-buckets (case 2), the objective is to balance the size of the two created super-buckets. To address case 2, a straightforward greedy strategy is employed to split the super-bucket into two smaller ones. We propose a simple greedy algorithm (see Algorithm 1) for performing this task.

#### Algorithm 1

Divide a super-bucket into two super-buckets

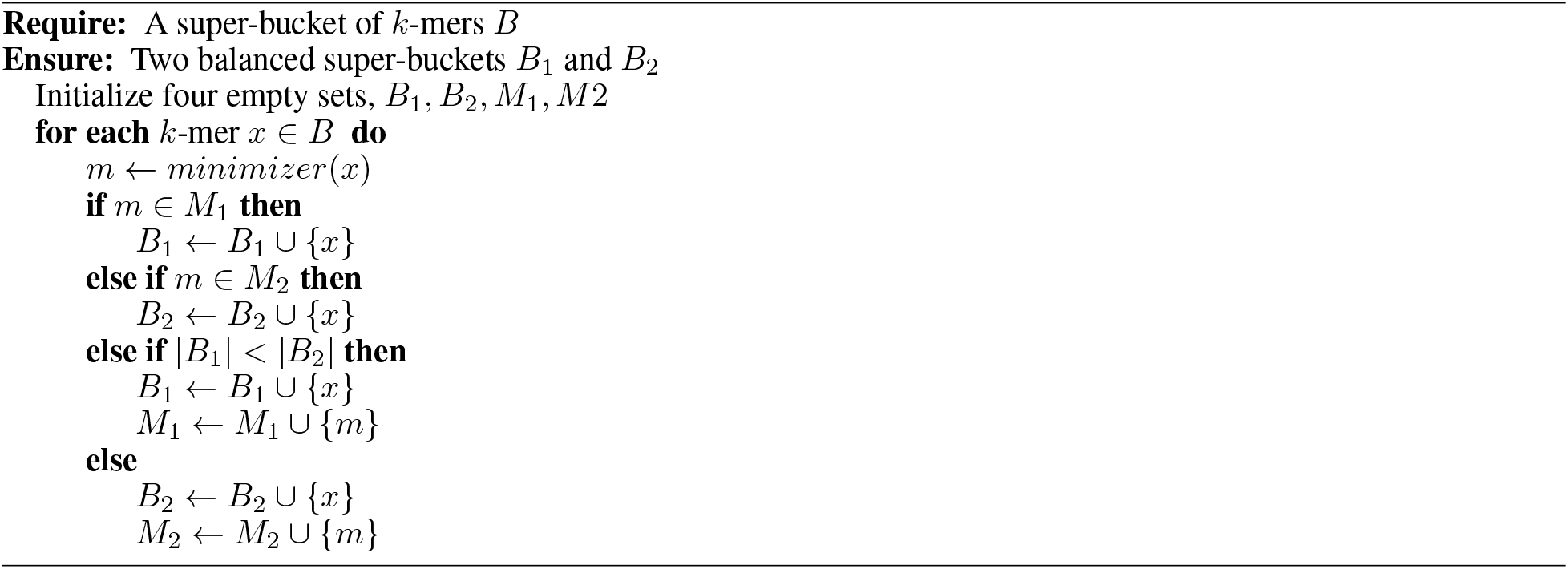

### 2.6 Read querying

A compacted de Bruijn graph constructed using `Cdbgtricks` supports sequence queries. In practice, while querying a sequence *s* on a graph *G*, `Cdbgtricks` determines if at least *α*% of the *k*-mers of *s* are in the graph (with *α* a user-defined parameter). If this is the case, `Cdbgtricks` indicates the uni-MEMs, as defined in `deBGA` [5]. Each uni-MEM is a tuple *< u*_*id*_, *u*_*start*_, *u*_*end*_, *s*_*start*_, *s*_*end*_ *>* where *u*_*start*_ and *u*_*end*_ are the start and end positions of mapping on the unitig whose identifier is *u*_*id*_, and *s*_*start*_ and *s*_*end*_ are the start and end positions of mapping on the queried sequence *s*. In other words, the *k*-mers whose offsets in the read are between *r*_*start*_ and *r*_*end*_ are found in the unitig *u*_*id*_ between *u*_*start*_ and *u*_*end*_. A uni-MEM is found through the extension of the first common *k*-mer between the read and a unitig. The extension ends in one of the following cases:

1. A mismatch is encountered.
2. Either the end of the read or the end of the unitig is encountered.

## 3 Results

All presented results are reproducible using command lines and versions of tested tools, that are given in this repository https://github.com/khodor14/Cdbgtricks_experiments. The executions were performed on the GenOuest platform on a node with 4 *×* 8 cores Xeon E5-2660 2,20 GHz with 128 GB of memory.

### 3.1 Genome datasets

We tested `Cdbgtricks` in two frameworks, corresponding to two input datasets of distinct size and complexity. The first one, called “*human*” is composed of 100 assembled human genomes that were used in the `GGCAT` experiments [14]. These genomes are available on zenodo (10.5281/zenodo.7506049,10.5281/zenodo.7506425). The second set, called “*coli*” is composed of 7055 *E. coli* genomes downloaded from NCBI (https://ftp.ncbi.nlm.nih.gov/genomes/all/GCF/030/).

While `Cdbgtricks` is capable of initially constructing a compacted de Bruijn graph from scratch, it is worth noting that there are faster alternatives for creating the initial graph. As such, for each dataset, we created an initial graph in fasta format from one genome (chosen as the first in the alphabetic order of the file names) using `Bifrost`. Once created `Cdbgtricks` can be used for indexing this initial graph. Subsequently, for each dataset, we added one by one the remaining genomes.

### 3.2 Used Parameters

The used parameters are the same for the two datasets. In all experiments we used *k* = 31 and minimizers of size *m* = 11. During the update experiments the tools were executed using 32 threads, while during the query experiments the tools were executed using a single thread.

For `Cdbgtricks`, the parameters controlling the bucket size were set to default. The bucket lower bound size is *ρ* = 5000 and a the super-bucket multiplicative factor *γ* is set to 4. This setting of parameters means that the size of a super-bucket is approximately 20000 *k*-mers, and once it reaches 40000 *k*-mers, it gets divided into two super-buckets. The values of the parameters were chosen to ensure satisfactory results that will be shown in the subsequent sections.

### 3.3 Percentage of modified buckets

As explained Section 2.3, one of the key ideas in `Cdbgtricks` is to distribute indexed *k*-mers into multiple buckets, each bucket being indexed with its own MPHF. Doing so, we expect that, while adding *k*-mers from novel sequences, *k*-mers are added to only a fraction of the buckets, and then only, a fraction of the MPHFs have to be recomputed. More precisely, we expect that the percentage of modified buckets, i.e. 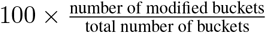 decreases as the number of genomes in the graph increases. Note that here we do not differentiate buckets and super-buckets and we regroup these two notions in the term “buckets”.

In this section we test this expectation on the *human* and *E. coli* datasets. The results about the percentage of modified buckets are shown Figure 5. Results show that, as expected, the percentage of modified buckets decreases with respect to the number of genomes. The shaded cluster of points for the *E. coli* dataset shows that in the majority of cases, the percentage of modified buckets and super-buckets is less than 20%. More specifically, results with, say, more than 5000 *E. coli* genomes show that, except for some outliers, less than 10% of the buckets are modified when adding a new genome.

**Figure 5.**
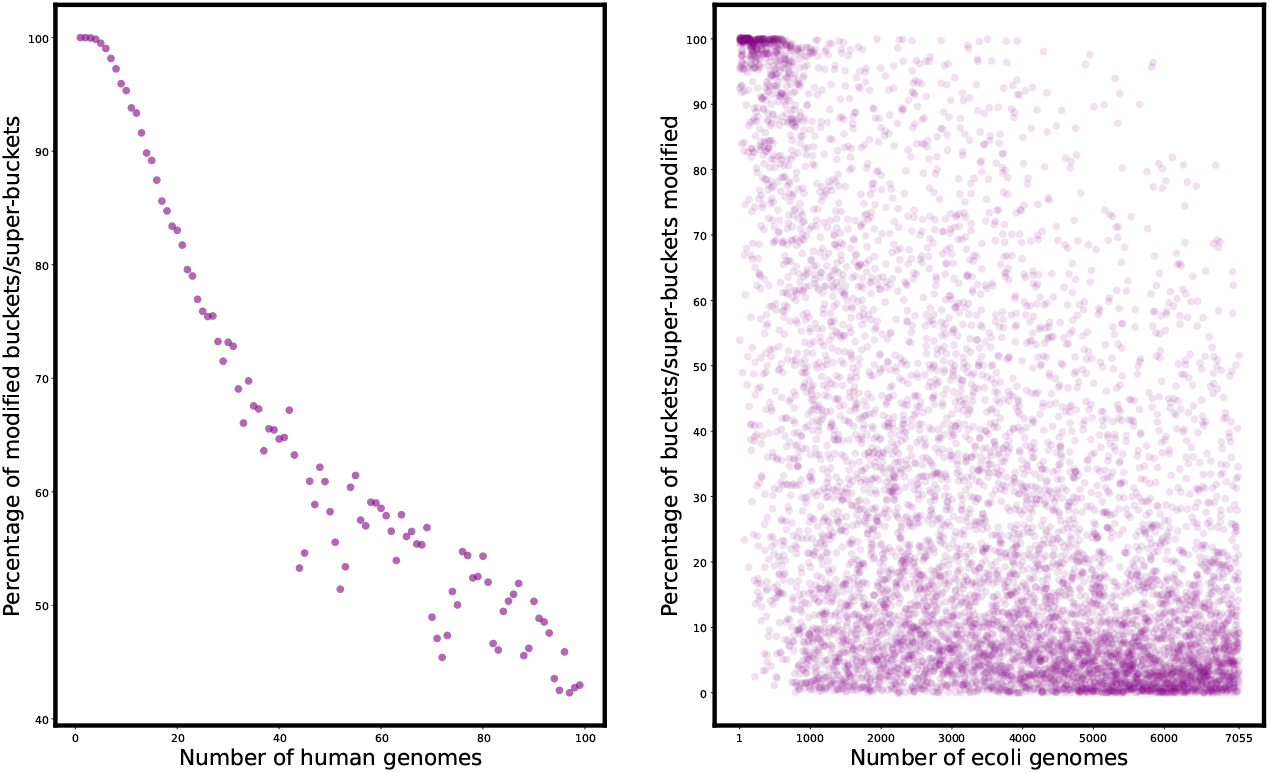
Percentage of modified buckets.

These results validate the chosen default parameters, and they confirm the expectation that lesser and lesser buckets are modified while increasing the number of genomes of the same species in a compacted de Bruijn graph.

### 3.4 Scalability

One of the main objective of `Cdbgtricks` is the time performances when updating a compacted de Bruijn graph with new sequences. In this context, we compared the `Cdbgtricks` update time, with the update time obtained thanks to `Bifrost`, also able to update an already created compacted de Bruijn graph. Furthermore, although `GGCAT` does not provide graph updating capabilities, we included it in our comparison due to its efficiency. The memory and disk for the update with `Cdbgtricks` and `Bifrost` and for the construction with `GGCAT` are also reported.

Results for the *human* dataset are shown in Figure 6. Note that, on the *human* dataset, `GGCAT` reached a timeout we set at two days on more than 71 human genomes. Hence, only the results for the first 71 genomes were reported for `GGCAT`. Globally, the results on this dataset show that `Cdbgtricks` is at least 2x faster than `Bifrost` on graphs composed of 50 genomes or more. Compared to `GGCAT`, `Cdbgtricks` is slower on this small number of genomes. Given that as the number of genomes in the graph increases, the `GGCAT` construction time naturally increases while the `Cdbgtricks` update time decreases, one can expect `Cdbgtricks` to be faster when dealing with more genomes than those tested here. However, given the observe `GGCAT` limitation after 71 genomes, we could not verify this fact in practice, at least for human genomes. The memory used by `Cdbgtricks` and `Bifrost` are slightly the same and are limited to a few dozen gigabytes. `GGCAT` uses much less memory, but needs up to order of magnitude more disk.

**Figure 6.**
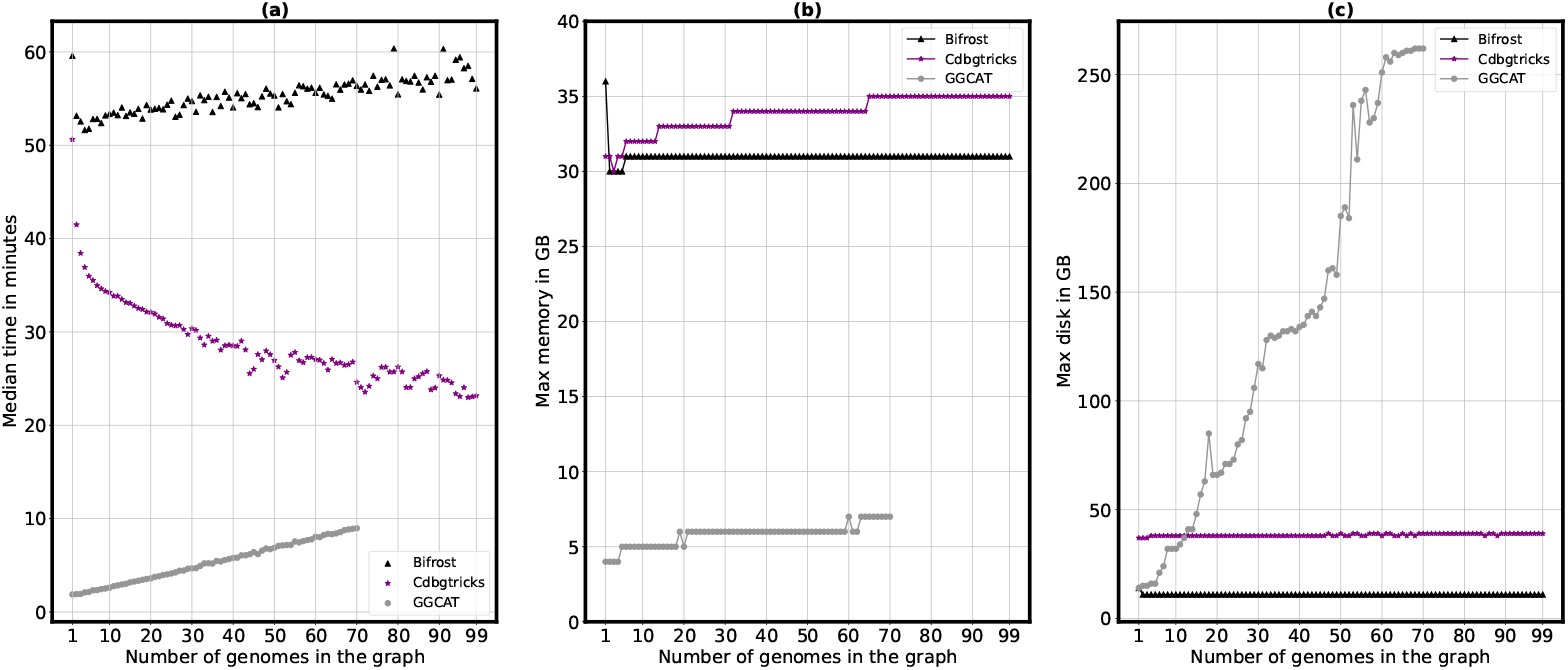
Results on human genomes dataset. Time (a), memory (b) and disk usage (c) are given for updating a graph for for `Cdbgtricks` and `Bifrost` and for constructing a graph from scratch for `GGCAT`.

Results for the *coli* dataset are shown in Figure 7. For the sake of clarity of presenting the results of the *E. coli*, we chose to report the median time over a window of 200 genomes. The detailed presentation of execution time is in supplementary materials. On this dataset, both `Bifrost` and `GGCAT` were faster than `Cdbgtricks` on graphs composed of less than a thousand genomes. However, in the vast majority of cases, when the number of genomes get higher than, say, 2000 genomes, `Cdbgtricks` is 2x to 3x faster than `GGCAT` and `Bifrost`. With `Cdbgtricks`, adding an *E. coli* genome to a compacted de Bruijn graph graph containing already few thousands genomes requires between 30 and 50 seconds. `Cdbgtricks` uses slightly the same amount of memory compared to `GGCAT` and roughly twice the amount of memory compared to `Bifrost`. `Cdbgtricks` uses up to 4x more disk compared to `Bifrost`, while it uses much less disk compared to `GGCAT`.

**Figure 7.**
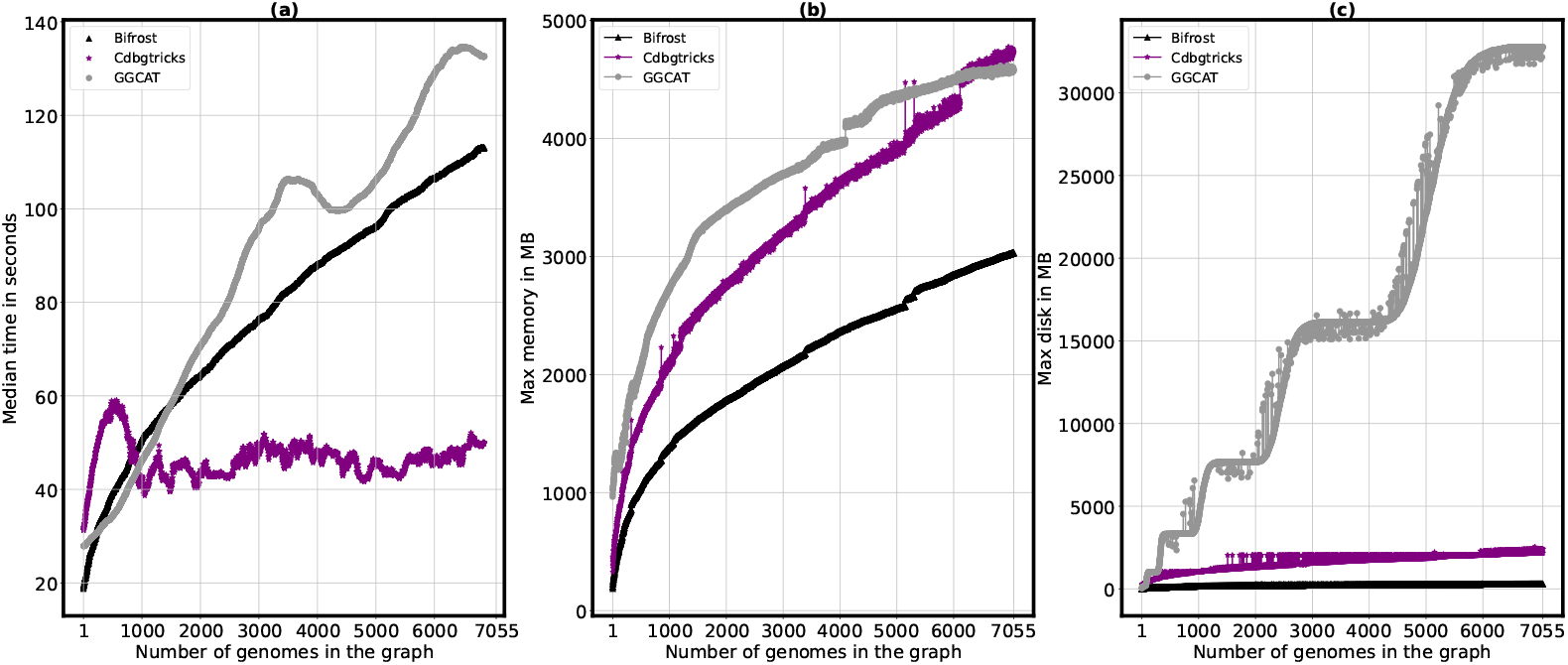
Results on *E. coli* genomes dataset. Time (a), memory (b) and disk usage (c) are given for updating a graph for for `Cdbgtricks` and `Bifrost` and for constructing a graph from scratch for `GGCAT`. The time is given as the median over a window of 200 consecutive points.

### 3.5 Results querying sequences

We propose some experiments for comparing the query performances of `Cdbgtricks` with those of `Bifrost`, `GGCAT` and `SSHash`. Note that these four tools do not offer the same query features. Although, these results must be considered as rough estimations showing the main tendencies.

Using `Bifrost`, we constructed a graph from 15,806 *E. coli* genomes, and a graph from 10 human genomes. Then we constructed an index for each graph using either `Cdbgtricks`, `Bifrost` or `SSHash`. We have differentiated between results obtained with positive queries (querying sequences present in the graph) and those obtained with negative queries (querying random sequences) with *k*-mers that are not in the two constructed graphs. The positive queries are a subset of unitigs from each graph. The negative queries are composed of one million random sequences of length between 500 and 1000 base pairs. The querying results are shown Table 1.

**Table 1:** Performances of sequence queries using a compacted de Bruijn graph for Cdbgtricks, Bifrost, SSHash, and GGCAT.

| Dataset | Query type | Tool | Memory (MB) | Disk (MB) | time (mm:ss) |
| --- | --- | --- | --- | --- | --- |
| *E. coli* | Negative | Cdbgtricks | 4723 | 0 | 10:02 |
|  |  | Bifrost | 4362 | 0 | 06:43 |
|  |  | SSHash | 725 | 0 | 00:07 |
|  |  | GGCAT | 560 | 3325 | 01:32 |
|  | Positive | Cdbgtricks | 4724 | 0 | 02:15 |
|  |  | Bifrost | 4362 | 0 | 01:43 |
|  |  | SSHash | 725 | 0 | 01:10 |
|  |  | GGCAT | 644 | 2978 | 01:26 |
| *human* | Negative | Cdbgtricks | 25520 | 0 | 12:25 |
|  |  | Bifrost | 27376 | 0 | 11:37 |
|  |  | SSHash | 6090 | 0 | 00:07 |
|  |  | GGCAT | 615 | 6861 | 4:55 |
|  | Positive | Cdbgtricks | 25520 | 0 | 04:23 |
|  |  | Bifrost | 27376 | 0 | 06:37 |
|  |  | SSHash | 6090 | 0 | 01:14 |
|  |  | GGCAT | 746 | 7053 | 05:04 |

The results shows that `Cdbgtricks` and `Bifrost` obtained similar results in term of memory, while `Cdbgtricks` is slightly slower. While `GGCAT` is faster than `Bifrost` and `Cdbgtricks`, and it uses the smallest amount of memory in these query experiments, it uses few Gigabytes of disk. Despite `SSHash` being the fastest tool, it consumes an order of magnitude more memory than `GGCAT`. These results shows that `Cdbgtricks` offer queries in a reasonable amount of time, and the performance of `Cdbgtricks` is close to the performance of `Bifrost`.

## 4 Discussion and future work

In this paper, we presented `Cdbgtricks`, a novel method for updating a compacted de Bruijn graph when adding new sequences such as full genomes. `Cdbgtricks` also indexes the graphs, hence it enables to query sequences and detect the portions of the graph that share *k*-mers with the query. The dynamicity of the proposed index is achieved thanks to the distribution of *k*-mers into multiple buckets, each bucket being indexed using a minimal perfect hash function (MPHF). The addition of new *k*-mers affects only a fraction of the buckets, for which the MPHF has to be recomputed. In practice, when indexing a large number of genomes (dozens of human genomes or thousand of *E. coli* genomes) `Cdbgtricks` outperforms the computation time of state-of-the-art tools dedicated to the creation of the update of compacted de Bruijn graphs.

Of independent interest, exploiting `PTHash`, the `Cdbgtricks` indexing framework offers a theoretical way to bijectively associate each *k*-mer from a set composed of *n* distinct *k*-mers with a unique value in [0, *n*[. In that respect, despite the fact that some engineering work remains to be done to achieve this practical feature, our indexing strategy can be used as an MPHF for this kind of dataset. Furthermore, it will present two main additional advantages when compared to a classical MPHF:

- In essence, an MPHF is static. Adding an element to this kind of data structure requires one to recompute the entire MPHF from scratch to associate the *n* + 1 elements to a unique value in [0, *n* + 1[. As `Cdbgtricks` distributes the *k*-mer set over numerous “sub-MPHFs”, adding an element requires only to recompute one of the sub-MPHFs. This offers a clear advantage when adding a few elements to large MPHFs, composed of, say, billions of elements.
- The MPHF definition does not impose that so-called “alien *k*-mers” (*k*-mers not belonging to the indexed set) are detected as aliens at query time. Actually, MPHFs that do not store the indexed set of elements (as this is the case for `BBhash` and `PTHash`), are not able to always discriminate an alien *k*-mer from an indexed one. In the context of this work, the presence of the indexed *k*-mers in the stored unitigs enables us to validate that a query *k*-mer actually belongs to the original set, and thus enables us to detect whether it is an alien *k*-mer or not.

A future research direction is to devise a smarter bucket clustering approach. One way could be to group the buckets whose minimizers appear in the same unitigs. Doing so, we could expect more data locality, limiting the cache-misses.

The compacted de Bruijn graph computed by `Cdbgtricks` is not colored. This means that the information is lost about the original genome(s) a *k*-mer belongs to. In recent years, significant attention has been given to the use of colored and compacted de Bruijn graphs in computational biology applications [24]. Hence, another research priorities for the future of this tool is to integrate the color information. This will necessitate minor yet potentially expensive operations. To include the new colors associated with a set of new sequences *S*, the color information of all *k*-mers in *S* already present in the graph *G* will have to be updated, while those *k*-mers are not modified in the current uncolored `Cdbgtricks` version.

There exists no limitation for using `Cdbgtricks` for merging the information of two compacted de Bruijn graphs *G*_1_ and *G*_2_. We can simply consider the *k*-mers of, say, *G*_2_ to be added into *G*_1_, and apply the exact same algorithm as proposed here. Future work will include validation and scaling tests for this approach.

Finally, we believe that the number of common *k*-mers but also the number of splits and joins performed when adding a sequence to a graph could be used as metrics to estimate the distance between a sequence and a compacted de Bruijn graph, or even between two compacted de Bruijn graphs.

## Acknowledgements

This project received funding from the European Union’s Horizon 2020 research and innovation program 369 under the Marie Skłodowska-Curie grant agreement No 956229. We acknowledge the GenOuest bioinformatics core facility https://www.genouest.org for providing the computing infrastructure.

## Notes

https://github.com/khodor14/Cdbgtricks

### Competing Interest Statement

The authors have declared no competing interest.

### Summary of Updates

The query section in the manuscript has been revised as well as the corresponding experiments.

