## Supplementary material for "Cdbgtricks: Strategies to update a compacted de Bruijn graph"

### Supplementary materials

#### 1 Figures

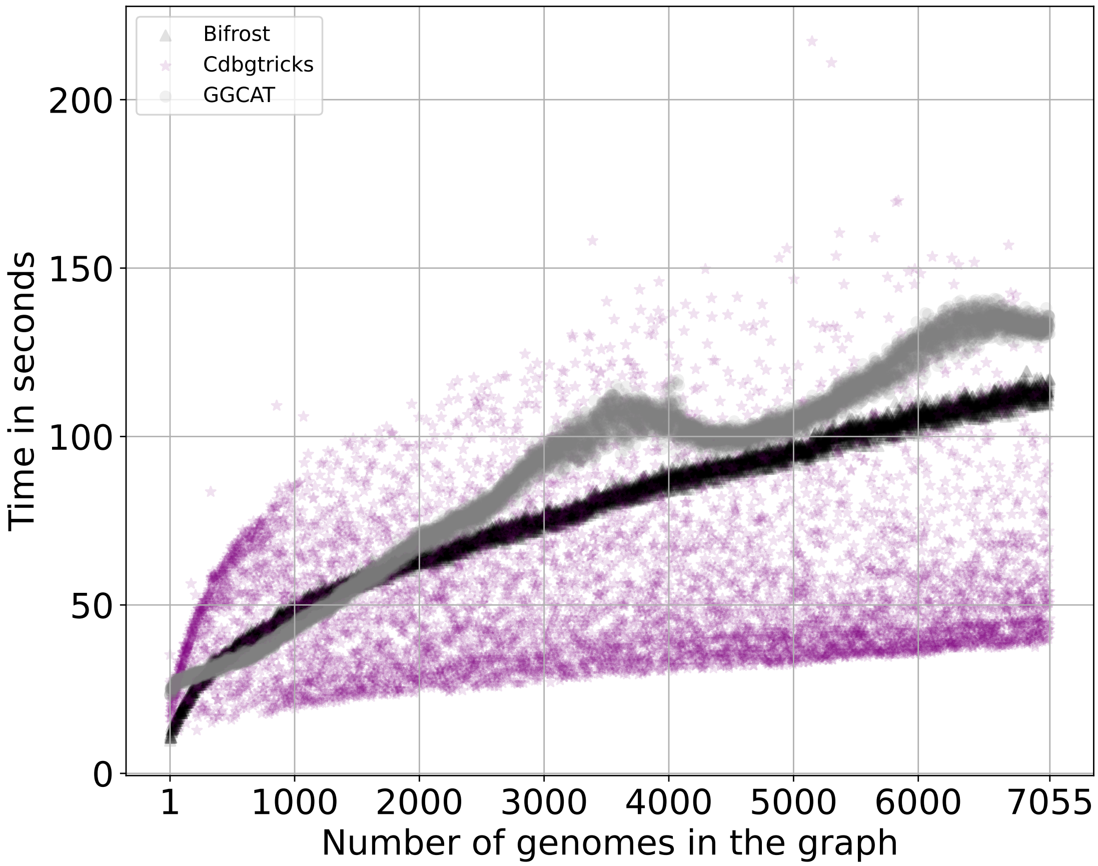


Figure 1: **Results on *E. coli* genomes dataset.** Time is given for updating a graph for for and and for constructing a graph from scratch for .

#### 2 Algorithms

**Algorithm 1** Test k-mer presence

**Require:** Hash Table of unitigs U, k-mer s, Table of buckets T, Table of mphfs F
**Ensure:** s belongs to a unitig

```
function KMERPRESENCE(H,s,T,F)
    mini_s ← canonicalminimizer(s)            ▷ compute the canonical minimizer of s
    i ← T[mini_s]                             ▷ retrieve the identifier of the bucket of mini_s
    if i ≠ NIL then                           ▷ the minimizer is in the index
        f_i ← F[i]                            ▷ retrieve the mphf of the bucket
        q ← canonical(s)                      ▷ compute the canonical form of s
        j ← f_i(q)                            ▷ compute the hash value of q
        p_j ← P[i][j]                         ▷ retrieve the position tuple of q
        id ← p_j.u_id                         ▷ retrieve the unitig identifier
        o ← p_j.u_off                         ▷ retrieve the offset of q in this unitig
        u ← U[id]                             ▷ retrieve the unitig
        return canonical(u(o, o + k)) = q     ▷ compare the k-mers
    end if
    return false                              ▷ the minimizer is not in the index, so s is not in the graph
end function
```

**Algorithm 2** Construct funitigs

```
Require: Hash table of new k-mers K,Index of the graph I
Ensure: A set of funitigs X
  function CONSTRUCTFUNITIG(K)
    X ← ∅
    for each k-mer s ∈ K do
      if s is not marked as used then
        mark s as used
        s_1 ← ExtendRight(s)
        s_2 ← ExtendRight(s̄)
        X ← X ∪ {s̄_2 ⊙ s_1(k − 1, |s_1|)}
      end if
    end for
    return X
  end function
  function EXTENDRIGHT(s)
    q ← s
    g ← s(1, k)                                   ▷ the (k-1)-suffix of s
    while True do
      a ← FindRightExtensions(g)
      if a ≠ ϵ then
        b ← FindRightExtensions(ḡ)
        if b ≠ ϵ then
          q ← q ⊙ a
          g ← g(1, k − 1) ⊙ a
        else
          break
        end if
      else
        break
      end if
    end while
    return q
  end function
  function FINDRIGHTEXTENSIONS(x)                  ▷ x is a (k-1)-mer
    extension ← ϵ
    count ← 0
    for each a ∈ ∑ do
      q ← x ⊙ a
      if q ∈ K then
        extensions ← a
        count ← count + 1
      else if KMERPRESENCE(I.U,s,I.T,I.F) then     ▷ x is found in the graph
        extension ← ϵ
        break
      end if
    end for
    if count ≠ 1 then
      extension ← ϵ
    end if
    return extension
  end function
```
